# Infra Red Thermography Reveals Transpirational Cooling in Pearl Millet (*Pennisetum glaucum*) Plants Under Heat Stress

**DOI:** 10.1101/2020.09.04.283283

**Authors:** Arun K. Shanker, Divya Bhanu, Basudev Sarkar, S.K. Yadav, N. Jyothilakshmi, M. Maheswari

**Affiliations:** ICAR-Central Research Institute for Agriculture Santoshnagar, Saidabad P.O, Hyderabad – 500059, India

**Keywords:** Infra red imaging, heat stress, thermography, high temperature, evaporative cooling

## Abstract

An infra red thermographic analysis of well watered control and well watered heat stressed pearl millet (*Pennisetum glaucum*) was conducted at ICAR – Central Research Institute for Dryland Agriculture as a part of high resolution phenomics studies to identify the individual quantitative physiological parameters by plant phenotyping that form the basis for more complex abiotic stress tolerant traits. It was seen that the temperature gradient increased gradually from ground level to the top in the control non heat stressed plant. In contrast, it was seen that in the heat stressed plant the temperature increased up to the middle of the plant and then started to decrease at the top of the plant in comparison with the non heat stressed control plant. Our results indicate that the lowering of temperatures in the top of the heat stressed plant may be a mechanism by which the heat stressed plant acclimates to stress by regulating its transpiration thereby bringing in a cooling effect to counter stress.

## Introduction

Temperature which is the mean kinetic energy of the molecules within a system measured in Kelvin or Celsius is a fundamental characteristic of the climate in an ecosystem. The main concern of global warming and the changing climate is effect of rising temperatures on plants and in turn their effects on growth and development and yield in the case of agricultural crops (Sun et al 2019). Temperature has a plethora of influences across a range of biological processes from enzyme bioenergetics to ecosystem biogeochemistry. It plays a central role in the primary control of processes at spatial and temporal scales from plants to ecosystems. Temperature gradients along the plant from the soil plant interface region towards to the apex of the plant can be of interest due to the subtle but significant control it exerts on various physiological, biochemical and molecular processes in the plant (Legris et al 2017). Leaf temperature specifically is of paramount importance as it is related to transpiration rate, stomatal conductance, vapor pressure deficit and boundary layer resistance at the leaf surface. Pearl millet (*Pennisetum glaucum*) is one the most widely grown millet in the world and has been prominent in continent of Africa and Asia since prehistoric times. It is highly tolerant to drought, heat and acid soils. The grain is highly nutritious and has a good value as a food grain in the tropics (Andrews and Kumar 1992; Varshney et al 2017).

Thermal imaging of plants at Infra Red spectrum between 9000 –14000 nm is a well studied method for detection of various abiotic and biotic stress in plants (Jones 2018; Prashar and Jones 2014). Recent advances in thermal imaging technology offers specific prospects to take up robust high resolution phenotyping in plants with emphasis on temperature gradient from collar region to the apex of the plant. These techniques help us conduct real time temperature screening for detection of physiological changes in the plants under different environmental conditions allowing us to monitor plants non-destructively. The microclimate near the leaf surface is affected by leaf temperature which in turn affects the physiological processes in the plant. Plants can have different strategies to maintain leaf temperature at optimum level so that the physiological process are not affected (Lin et al 2017). Thermal imaging can be a non destructive tool for screening for drought tolerance in plants (Biju et al 2018). Temperature measurements and thermography can be used in many ways to address physiological questions such as the extant of control temperature can have on the plant processes (Still et al 2019).

We hypothesized that temperature differences in the plant can accompany specific strategies employed by plants, such as structural and functional changes in the water uptake and transpiration systems by which they acquire new homeostasis which may aid in protective adaptations. The changes observed in the plants under stress may be as a response to stimuli or alterations to counter stress. We conducted an experiment with the objective to study the individual quantitative physiological parameters by plant phenotyping that form the basis for more complex abiotic stress tolerant traits.

## Materials and Methods

We conducted an experiment in ICAR – Central Research Institute for Dryland Agriculture (ICAR-CRIDA) in July – August 2020 on pearl millet (*Pennisetum glaucum*). Two varieties of Pearl millet - ICMH 356 and 86M86 was used for the study obtained from International Crops Research Institute for the Semi-Arid Tropics (ICRISAT), Hyderabad. The variety ICMH is developed from parents - ICMA 88004 x ICMR 356 ICRISAT 1993, has a duration of 75-80 days, is medium tall and semi-compact with thick conical earheads, yellow anthers, obovate bold yellow brown grains.

It has a mean yield of 2.3-2.5 T /ha. The variety 86M86 is from Parents - M128F x M138R, it has a good yield performance in different soil conditions, high capability of grain production, strong stem with strong root system and resistant to lodging, stay green till maturity, ready to harvest in approximately 95-100 days (after seed emergence). A total of 72 pots were used for growing pearl millets in open atmospheric conditions with a mean day length of 12.5 hours. The treatments were T1 – Control, T2 – Water stress, T3 – Heat stress and T4 – Water + Heat Stress. There were 7 replications of each treatment and two sets of experiments one for each variety was maintained.

The pots were irrigated every with tap water. Water stress treatment was imposed on water stress and Water stress + Heat treatments at 43 DAS by withholding irrigation. Heat treatment was imposed so as to coincide with water stress treated plants. Heat treatment was imposed on 48 DAS plant in heat and heat + water stress treatments by transferring the pots to green house conditions where the ambient temperature was 3 degrees more than the ambient temperature in the open. The mean high temperature in open conditions was 28.7 °C and 22.1 °C, the heat treatment had mean high temperature of 34.1 °C and a minimum of 32.7 °C. After 5 days of treatment, thermal imaging of the plants was done with Infra Red camera FLIR E-95 with laser-assisted autofocus and on-screen area measurement, 161,472 (464 × 348) points of temperature measurement and wide temperature ranges. Images were taken at 4.9 meters distance from the plants at a relative humidity of 50 percent, the emissivity was at 0.95.

A soil moisture probe with mounted shaft TDR 350 - Field-Scout, Spectrum Technologies Inc., Aurora, USA was used to measure soil moisture content at different soil depths. The soil moisture probe was calibrated gravimetrically, raw sensor readings were compared with Volumetric Water Content (VWC) estimated by gravimetrically to calibrate the instrument. Soil moisture in terms of VWC was measured at 7.6, 12 and 20 cm depth of soil in the pots.

## Results and Discussion

Here we present the results of thermal images of control and heat stress treated plants both of which are well watered and discuss the implications in water relations as seen from temperature gradients observed in the plants from the collar region to the apex of the plants. The soil moisture content in terms of VWC at the after 5 days of imposition of stress is shown in Fig. 1. The VWC was 42.3 in control at 7.6 cm soil depth, 39.7 at 12 cm depth and 37.2 at 20 cm depth. In the Heat stressed pots the VWC was 34.6 at 7.6 cm soil depth, 32.7 at 12 cm soil depth and 31.9 at 20 cm soil depth. The Infra red image of the control plant and the heat stressed plant is shown in Fig. 2.

**Fig. 1.**
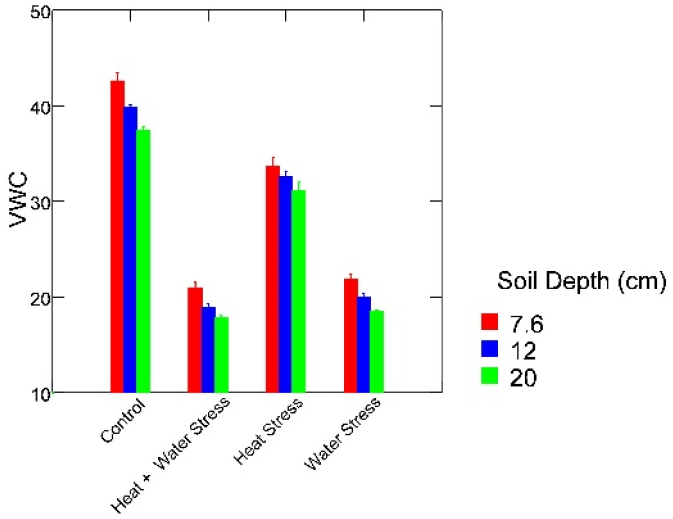
Soil moisture VWC at different soil depths in control and stress treatments

**Fig. 2.**
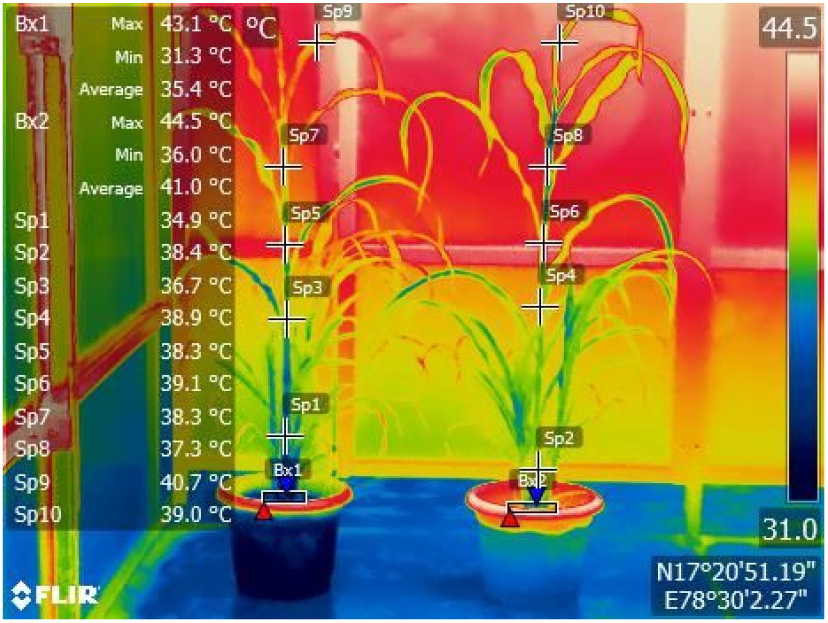
Infra Red Thermographic image of control and heat stressed pearl millet plants

The plant at the left is the control and the plant in the right is the heat stressed plant. The image shows the temperature gradient in the plants as measured by FLIR tools software which is used to analyze images acquired by FLIR E-95 camera. Box 1 is the soil plant interface region of the control plant where the temperature is Max 43.1 °C and Min was 31.3 °C, the mean temperature of the region was 35.4 °C. Box 2 is the soil plant interface region of heat stressed plant where the temperature is Max 44.5 °C and the Min was 36.0 °C and the mean temperature of the region covered by the box was 41.0 °C.

This was clearly indicative of the high water loss in the soil due to heat in the heat stressed plant which is in confirmation with Liu and Huang (2005). Spots 1, 3, 5, 7 and 9 are temperature gradient in the stem as we go up towards the apical region of the control plant. Spots 2, 4, 6, 8 and 10 are temperature gradient in the stem as we go up towards the apical region of the heat stressed plant. The temperature gradient in the control plants is 34.9 °C, 36.7 °C, 38.3 °C, 38.3 ° C and 40.7 °C at the top of the canopy. The temperature gradient in the heat stressed plant is 38.4 °C, 38.9 °C, 39.1 °C, 37.3 °C and 39.0 °C at the top of the canopy. A distinct blue colour is seen in the region of the stem between Spot 4 and Spot 8 which is not seen in the control plant indicating that the heat stressed plant it is at lower temperature in the middle region as compared to the control plant.

The temperature gradient is at a constant increase from ground level to the top in the control plant reaching a maximum of 40.7 °C at the top which may be due to high transpiration rate in the plant.

In contrast, it is seen that in the heat stressed plant the temperature increases up to Spot 6 in the middle of the plant and then starts to decrease as we go to the top of the plant. This may be due to evaporative cooling taking place in the plant as a result of higher transpiration rate due to heat. Thermography accurately estimates temperature of plant when imaged with IR cameras, the surface temperature of crop canopies tends to lower with increasing transpiration which is the result of cooling due to evaporation (Prashar and Jones 2014). The lower than control temperatures in the top regions of the plant canopy observed in the heat stressed plants may be because there is an increase in transpiration and in turn in cooling of the leaf surface reflecting in lower temperatures, there can be a decrease in leaf air vapour pressure deficit. This may be a mechanism by which the heat stress plant acclimates to stress by increasing its transpirational regulation thereby bringing in a cooling effect to counter stress (Lapidot et al 2019; Buckley et al 2019). It is seen from thermographic analysis of the control and heat stressed plants that evaporative cooling is an adaptive mechanism of the plant when exposed to extremes of temperature. Similar results has been reported (Crawford et al 2012) where high temperature exposure increased cooling. It is known that temporary storage of heat has a cooling effect (Seginer 1994).

There is an increase in possibility of heat damage at high temperatures which can result in water loss from the plant, this can be minimized by cooling of the leaf surface by water evaporation from the stomata. It is possible that in the well watered conditions the plant is transpiring at an optimum rate wherein cooling of the leaf surface is not a requirement, on the other hand under well watered heat stressed the plant is tending to transpire more inorder to reduce the leaf temperature as an adaptive mechanism.

## Supporting information

Supplementary Temperature Analysis Data

## Acknowledgements

The authors wish to thank the Director CRIDA for providing the facilities for the work.

The authors have no conflict of interest

